# Trapping of Nicotinic Acetylcholine Receptor Ligands Assayed by *in vitro* Cellular Studies and *in vivo* PET Imaging

**DOI:** 10.1101/2021.12.08.471775

**Authors:** Hannah J. Zhang, Matthew Zammit, Chien-Min Kao, Anitha P Govind, Samuel Mitchell, Nathanial Holderman, Mohammed Bhuiyan, Richard Freifelder, Xiaoxi Zhuang, Jogeshwar Mukherjee, Chin-Tu Chen, William N. Green

## Abstract

A question relevant to nicotine addiction is how nicotine and other nicotinic receptor membrane-permeant ligands, such as the anti-smoking drug varenicline (Chantix), distribute in brain. Ligands, like varenicline, with high pKa and high-affinity for α4β2-type nicotinic receptors (α4β2Rs) are trapped in intracellular acidic vesicles containing α4β2Rs *in vitro*. Nicotine, with lower pKa and α4β2R affinity, is not trapped. Here, we extend our results by imaging nicotinic PET ligands *in vivo* in mouse brain and identifying the trapping brain organelle *in vitro* as Golgi satellites (GSats). Two PET ^18^F-labelled imaging ligands were chosen: [^18^F]2-FA85380 (2-FA) with varenicline-like pKa and affinity and [^18^F]Nifene with nicotine-like pKa and affinity. [^18^F]2-FA PET-imaging kinetics were very slow consistent with 2-FA trapping in α4β2R-containing GSats. In contrast, [^18^F]Nifene kinetics were rapid, consistent with its binding to α4β2Rs but no trapping. Specific [^18^F]2-FA and [^18^F]Nifene signals were eliminated in β2 subunit knockout mice or by acute nicotine injections demonstrating binding to sites on β2-containing receptors. Chloroquine, which dissipates GSat pH gradients, reduced [^18^F]2-FA distributions while having little effect on [^18^F]Nifene distributions *in vivo* consistent with only [^18^F]2-FA trapping in GSats. These results are further supported by *in vitro* findings where dissipation of GSat pH gradients blocks 2-FA trapping in GSats without affecting Nifene. By combining *in vitro* and *in vivo* imaging, we mapped both the brain-wide and subcellular distributions of weak-base nicotinic receptor ligands. We conclude that ligands, such as varenicline, are trapped in neurons in α4β2R-containing GSats, which results in very slow release long after nicotine is gone after smoking.

**Significance:** Mechanisms of nicotine addiction remain poorly understood. An earlier study using *in vitro* methods found that the anti-smoking nicotinic ligand, varenicline (Chantix) was trapped in α4β2R-containing acidic vesicles. Using a fluorescent labeled high-affinity nicotinic ligand, this study provided evidence that these intracellular acidic vesicles were α4β2R-containing Golgi satellites. *In vivo* PET imaging with F-18 labeled nicotinic ligands provided additional evidence that differences in PET ligand trapping in acidic vesicles were the cause of differences in PET ligand kinetics and subcellular distributions. These findings combining *in vitro* and *in vivo* imaging revealed new mechanistic insights into the kinetics of weak base PET imaging ligands and the subcellular mechanisms underlying nicotine addiction.

## Introduction

Tobacco use continues world-wide and is the leading cause of preventable deaths in the United States (U.S. Department of Health and Human Services 2014). Nicotine, the addictive molecule in tobacco, binds to high-affinity nicotinic acetylcholine receptors (nAChRs) in the brain, where it initiates its addictive effects. nAChRs are members of the Cys-loop family of ligand-gated ion channels, all of which are pentameric neurotransmitter receptors (Karlin 2002, Albuquerque, Pereira et al. 2009). In mammalian brains, high-affinity nicotine binding sites are largely nAChRs containing α4 and β2 subunits (α4β2Rs) (Albuquerque, Pereira et al. 2009). In mice, knockout of either subunit reduced the pharmacological and behavioral effects of nicotine (Picciotto, Zoli et al. 1998, Marubio, Gardier et al. 2003), and this effect can be rescued by targeted β2 subunit expression in the mid-brain reward ventral tegmental area (VTA); (Maskos, Molles et al. 2005).

Chronic nicotine exposure causes α4β2R upregulation, a complex set of long-term changes causing both increases in α4β2R high-affinity binding site numbers and functional upregulation, the increased α4β2R functional response after nicotine exposure (Marks, Burch et al. 1983, Schwartz and Kellar 1983, Benwell, Balfour et al. 1988, Breese, Adams et al. 1997). Several other studies link the process of nicotine-induced α4β2R upregulation to nicotine addiction (Vezina, McGehee et al. 2007, Govind, Vezina et al. 2009, Govind, Walsh et al. 2012, Lewis and Picciotto 2013). Nicotine and other weak-base ligands of α4β2Rs, unlike acetylcholine, are highly membrane-permeant and rapidly reach equilibrium in intracellular organelles and concentrate in acidic organelles (Brown and Garthwaite 1979). The anti-smoking drug, varenicline (Chantix), is selectively trapped as a weak base within intracellular acidic vesicles that contain high-affinity α4β2Rs because of its high pKa and binding-affinity (Govind, Vallejo et al. 2017). Nicotine, with lower pKa and affinity, is not trapped but concentrates in vesicles. Nicotine upregulation amplifies varenicline trapping by increasing the number of high-affinity binding sites, which increases the capacity of the acidic vesicles to trap varenicline and increasing acidic vesicle numbers.

Nicotine and varenicline bind to the same site on α4β2Rs in the brain. However, nicotine residence time in the brain is 1-2 hours while that for varenicline is 4-5 days, which may play a significant role in understanding its smoking cessation properties and the development of new cessation strategies (Govind, Vallejo et al. 2017). Positron emission tomography (PET) is an imaging modality that monitors ligand binding to nicotinic receptors *in vivo* (Mukherjee, Lao et al. 2018) and is, thus, an assay that could be used to examine differences between α4β2R ligands *in vivo* in the brain. PET ligands 2-[^18^F]FA85380 (2-FA) and [^18^F]Nifene were developed as nicotine analogs to monitor α4β2Rs *in vivo*. The fast *in vivo* kinetics of [^18^F]Nifene primarily resembles nicotine, while the slow kinetics of 2-[^18^F]FA closely resembles varenicline, suggesting that these tracers can be used to study the mechanisms of addiction and smoking cessation *in vivo*.

Here we address whether the distribution of PET tracers used to assay α4β2Rs *in vivo*, which are weak bases, are affected by trapping in α4β2R-containing acidic vesicles. Because of the limits of PET image resolution, we performed *in vitro* cell binding competition assays to assess ligand trapping, which allow subcellular resolution using the same PET ligands. We present evidence that 2-FA used as a PET tracer is trapped in α4β2R-containing acidic vesicles similar to varenicline trapping while Nifene is not trapped similar to nicotine. The identity of α4β2R-containing acidic vesicles is unknown. Recently, we found that a novel intracellular compartment in neurons, Golgi satellites (GSats), contain high levels of α4β2Rs and GSat numbers are increased by nicotine exposure as well as increased neuronal activity (Govind, Jeyifous et al. 2021). Using a fluorescent-tagged α4β2R ligand, Nifrorhodamine, to image the subcellular localization of α4β2Rs, we present evidence that α4β2Rs accumulate in acidic vesicles that are GSats. Live PET imaging in mice using [^18^F]Nifene and 2-[^18^F]FA were consistent with *in vitro* results with the kinetics of 2-[^18^F]FA PET signals were much slower than that of [^18^F]Nifene consistent with 2-[^18^F]FA being trapped in acidic vesicles. Chloroquine, which dissipates the pH gradient across acidic organelles, reduced trapping *in vitro* and reduced 2-[^18^F]FA’s distribution volume ratios (DVR) in midbrain and thalamus while having little to no effect on [^18^F]Nifene DVR values. Altogether, we conclude that 2-FA is trapped and images α4β2Rs in Golgi satellites while Nifene images all ligand-binding α4β2Rs because it is not confined to Golgi satellites.

## Material and Methods

### Cellular Binding Competition Assay

HEK cells stably expressing α4β2Rs were treated with 10 μM nicotine overnight. Cells were then washed four times with PBS, scraped off the plates, and suspended in PBS. Aliquots of cells were incubated with indicated concentrations of 2-FA, Nifene or nicotine respectively for 5 min followed by addition of 2.5 nM [^125^I]-epibatidine ([^125^I]Epb) (2200 Ci/mmol; Perkin Elmer) and incubation for 20 min at room temperature. At the end of incubation, cells were harvested on Whatman GF/B filters presoaked in 0.5% polyethyleneimine and washed four times with PBS using a 24-channel cell harvester (Brandel, MD). Non-specific binding was estimated by incubating parallel samples in 1 mM nicotine with [^125^I]Epb. Radioactivity of bound [^125^I]Epb was determined using a gamma counter (Wallac, Perkin Elmer, MA). The binding of [^125^I]Epb was normalized to the binding of cells without 2-FA, Nifene or nicotine treatment and plotted as percent of untreated cells.

### Mammalian cell culture

Stable cell lines expressing rat α4β2 nAChRs in HEK cells are from our lab, expressing untagged α4 and C-terminal, HA epitope-tagged β2 subunits (Vallejo, Buisson et al. 2005). Cell lines were maintained in DMEM (Gibco, Life technologies) with 10% calf serum (Hyclone, GE Healthcare Life Sciences, UT) at 37°C in the presence of 5% CO_2_. DMEM was supplemented with Hygromycin (Thermofisher scientific, MA) at 0.4 mg/ml for maintaining selection of α4β2 HEK cells. Stable cells were plated in media without hygromycin for experiments.

### Primary neuronal culture and transfections

Primary cultures of rat cortical neurons were prepared as described (Govind, Walsh et al. 2012) using Neurobasal Media (NBM), 2% (v/v) B27, and 2 mM L-glutamine (all from Thermofisher Scientific, MA). Dissociated cortical cells from E18 Sprague Dawley rat pups were plated on slips or plates coated with poly-D-lysine (Sigma, MO). For live imaging, neurons were plated in 35 mm glass bottom dishes (MatTek, MA). Cells were plated at a density of 0.25X10^6^ cells/mL on 35 mm dishes or per well in a 6-well plate. Neuronal cultures were transfected at DIV 10 with the Lipofectamine 2000 transfection reagent (Thermofisher Scientific, MA) according to manufacturer’s recommendations. Neurons were transfected with cDNAs of α4, β2_HA_ and Golgi markers, St3 GFP or St3 Halo. 0.5 μg of each DNA up to a total of 1.5 μg were used per 35 mm imaging dish or per well of a 6-well plate. 24 hours after transfection, neurons were treated with 1 μM nicotine for 17 hrs.

### Ligand treatment and ^125^I-epibatidine binding

α4β2R HEK cells were treated with 10 μM nicotine, 100 μM Nifene, or 100 μM 2-FA for 17 hours at 37°C in medium. Information for each ligand is displayed in Table 1. Cells were washed four times with PBS and collected by scraping off from the plates using a cell scraper with PBS, resuspended in 1ml PBS. 1/20^th^ of the cell suspension was distributed to three tubes followed by incubation in 2.5 nM [^125^I]-Epb for 15 minutes. For competition experiments, intact cells were preincubated in the indicated concentrations of Nifene, 2-FA or nicotine for 5 min, followed by the addition of 2.5 nM [^125^I]-Epb for 15 min. All binding was terminated by vacuum filtration through Whatman (Clifton, NJ) GF/B filters presoaked in 0.5% polyethyleneimine using a Brandel (Gaithersburg, MD) 24 channel cell harvester. Bound [^125^I]-Epb (2200 Ci/mmol) was determined by γ counting (PerkinElmer Wallac, Gaithersburg, MD) with nonspecific binding estimated by [^125^I]-Epb binding to cells that were preincubated with 1mM nicotine prior to epibatidine binding.

**Table 1.**
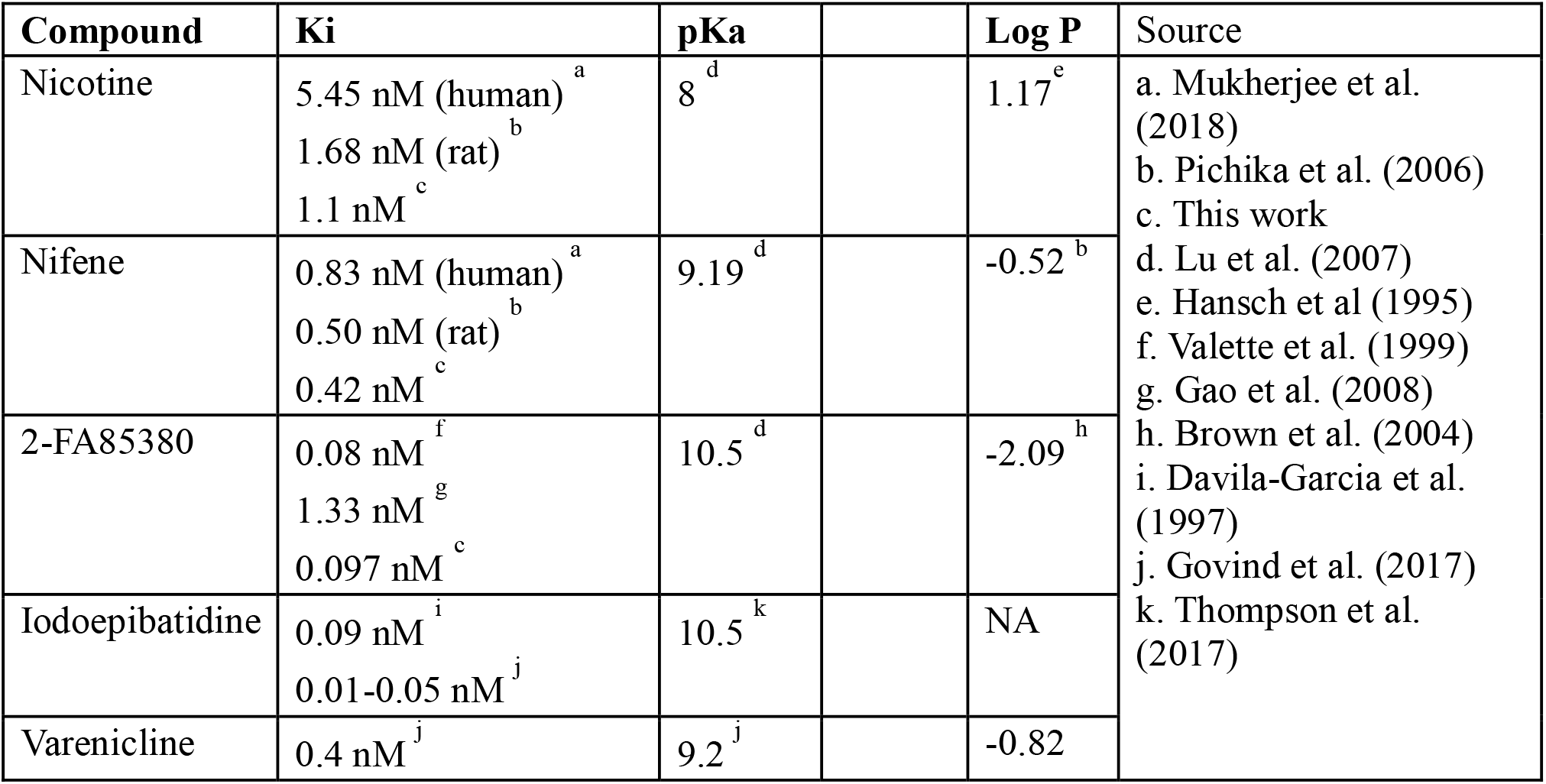
Nicotinic ligand acidity and affinity for α4β2 receptors.

### Altering intracellular pH of acidic intracellular organelles

Intact cells were treated with nAChR ligands for 17 hours and washed twice with PBS. Cells were exposed to two 5 min incubations with PBS containing 15, 150 μM chloroquine or 20 mM NH_4_Cl to neutralize the pH of intracellular organelles and washed two times with PBS. Cells were gently scraped from the plates, re-suspended in PBS and subjected to [^125^I]-Epb binding as described above.

### Immunocytochemistry

Live labeling experiments were performed on cultured neurons grown in imaging chambers (MaTek), transfected with nAChR subunits α4 and β2 and Golgi markers, St3GFP or St3 Halo for 24 hours and treated with 1 μM nicotine for 17 hours. Neurons were labeled with 25 μM Nifrorhodamine (analog of Nifrofam, (Samra, Intskirveli et al. 2018)) or pHRodo (Thermofisher) as per manufacturers instruction for 30 minutes, washed three times with low fluorescence Hibernate E buffer (Brain Bits, IL) and imaged live in the same buffer. For total β2 staining, neurons were grown on coverslips, fixed with 4% paraformaldehyde + 4% sucrose for 10 minutes and permeabilized with 0.1% triton X100. β2 subunits were imaged using monoclonal anti-β2 antibody, mAb12H and fluorescent secondary antibody, previously validated (Walsh, Roh et al. 2018), which gave results very similar to using mouse anti-HA antibody (Govind, Jeyifous et al. 2021). Primary antibody incubations were for 1 hour and secondary antibodies for 45 minutes unless otherwise mentioned. Coverslips were mounted using Prolong Gold mounting media with DAPI. Fluorescence images were acquired using a Marianas Yokogawa type spinning disk confocal microscope with back-thinned EMCCD camera. Images were processed and analyzed using ImageJ/Fiji (US National Institutes of Health).

### Animals

The male and female β2 nAChR knockout mice (KO) and their wild type (WT) littermates were generated in house by breeding a heterozygous pair on the C57BL/6J background purchased from the Jackson Lab (Bar Harbor, ME) (Picciotto, Zoli et al. 1995). Additional male and female WT mice at the same age were purchased directly from the Jackson lab and used in the same manner as the WT littermates. Animals were housed in The University of Chicago Animal Research Resources Center. The Institutional Animal Care and Use Committee of the University of Chicago, in accordance with National Institutes of Health guidelines, approved all animal procedures. Mice were maintained at 22–24°C on a 12:12-h light-dark cycle and provided food (standard mouse chow) and water ad libitum. Mice aged between 3 and 10 months were used in this study. The number of the animals and their body weights are summarized in Table 2

**Table 2.** Baseline information of the animals imaged with PET.

|  |  | WT | KO | AN | CQ |
| --- | --- | --- | --- | --- | --- |
| Sample, n | 2-[<sup>18</sup>F]-A85380 | 10 | 11 | 10 | 6 |
|  | [<sup>18</sup>F]-Nifene | 12 | 7 | 4 | 8 |
| BW, g | Male | 30.5 ± 0.6 | 30.6 ± 0.8 | 27.2 ± 2.2 | 29.9 ± 0.8 |
|  | Female | 21.6 ± 0.7 | 21.0 ± 0.5 | 18.8 ± 0.8 | 22.1 ± 0.4 |
| Injected dose of 2-[<sup>18</sup>F]-A85380 (μCi) |  | 192 ± 13 | 149 ± 13 | 201 ± 11 | 137 ± 7 |
| Injected dose of [<sup>18</sup>F]-Nifene (μCi) |  | 126 ± 7 | 87 ± 13 | 152 ± 2 | 146 ± 5 |

### Acute Nicotine Supplement

To demonstrate the binding specificity of weak base ligands, acute nicotine (AN) administration was performed to a group of WT mice. Nicotine ditartarte (Sigma-Aldrich, St Louis, MO) was prepared in isotonic saline at the concentration of 0.1 mg/ml freshly and filtered through a 0.2 μm syringe filter. Each mouse was injected 0.5 mg/kg body weight (BW) of nicotine intraperitoneally 15 min before the tracer administration and PET imaging according to the imaging protocol describe in the following sections.

### Chloroquine Treatment

Chloroquine diphosphate (CQ) (Sigma-Aldrich, St Louis, MO) was dissolved in PBS and filtered through a 0.2 μm syringe filter. WT mice were used in the CQ treatment group (CQWT). Each mouse received 50 mg/kg BW/day of CQ by intraperitoneal injection for 3 days (Vodicka, Lim et al. 2014). Ten minutes after the third CQ injection, animals underwent 2-[^18^F]FA or [^18^F]Nifene administration and PET imaging according to the imaging protocol described in following sections.

### Radiotracer Syntheses

Syntheses of both 2-[^18^F]FA and [^18^F]Nifene, two radiotracers for α4β2Rs, were carried out at the Cyclotron Facility of The University of Chicago. 2-[^18^F]FA was synthesized from the commercially available precursor, 2-TMA-A85380. The [^18^F]Nifene was synthesized from the precursor N-BOC-nitroNifene. An IBA Synthera V2 automatic synthesis module equipped with Synthera preparative HPLC was used for the radiolabeling and purification inside a Comecer hot cell. Typical yield for 2-[^18^F]FA was 34% (decay corrected) with specific activities >3,000 mCi/μmole and radiochemical purity > 99%. The representative radiochemical yield was 6.3% (decay corrected) with specific activities >3,000 mCi/μmole and > 99% purity for [^18^F]Nifene.

### PET/CT Imaging

The imaging protocols were designed based on previous reports for 2-[^18^F]FA and [^18^F]Nifene (Horti, Chefer et al. 2000, Pichika, Easwaramoorthy et al. 2006, Constantinescu, Garcia et al. 2013) and our preliminary experiments fine-tuning the protocol (data not shown). An intraperitoneal (IP) catheter was placed at the lower right abdominal area of each mouse before the imaging. The animal was then placed into the β-Cube preclinical microPET imaging system (Molecubes, Gent, Belgium) in a small animal holder. The tracer was delivered in 200 μL isotonic saline via the IP catheter at the same time when the PET imaging starts and followed by flushing 100 μL of fresh saline. The injected doses were summarized in Table 2. Whole body imaging was acquired with 133 mm x 72 mm field of view (FOV) and an average spatial resolution of 1.1 mm at the center of FOV (Krishnamoorthy, Blankemeyer et al. 2018). List-mode data was recorded for three hours for 2-[^18^F]FA and one hour for [^18^F]Nifene followed by a reference CT image on the X-Cube preclinical microCT imaging system, (Molecubes, Gent, Belgium). The images were reconstructed using an OSEM reconstruction algorithm that corrected for attenuation, randoms and scatter with an isotropic voxel size of 400 μm and a re-binned frame rate of 18 x10 minutes for 2-[^18^F]FA and 6 x10 minutes for [^18^F]Nifene. CT images were reconstructed with a 200 μm isotropic voxel size and used for anatomic co-registration, scatter correction and attenuation correction. Animals were maintained under 1-2% isoflurane anesthesia in oxygen during imaging. Respiration and temperature were constantly monitored and maintained using the Molecubes monitoring interface, and Small Animal Instruments (SAII Inc, Stoney Brook, NY) set up. All animals survived the imaging.

### Data Analyses of Tracer Uptake

To quantify the brain tracer uptake, analyses of CT-fused PET images were performed using VivoQuant software (Invicro, Boston, MA). Dynamic tracer uptake was analyzed and expressed in standardized uptake values (SUVs) in multiple 10-minute frames to generate time active curves (TACs) (Keyes 1995, Huang 2000, Hesterman, Ghayoor et al. 2019):

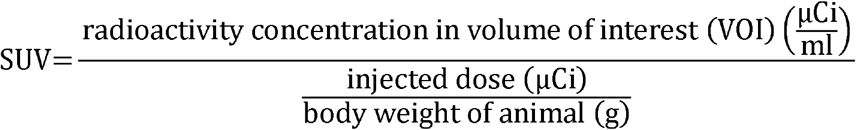

Brain regional analysis was performed using a 3-dimensional mouse brain atlas available as a VivoQuant software plug-in. The 3D brain atlas is based on the Paxinos-Franklin atlas registered to a series of high-resolution magnetic resonance images with 100 μm near isotropic data and has been applied in other studies (Slavine, Kulkarni et al. 2017, Hesterman, Ghayoor et al. 2019, Mazur, Powers et al. 2019). The brain was segmented into 14 regions within the skull of each animal using the CT as the reference for automatic registration. Among the 14 regions, 6 regions (cerebellum, cortex, hippocampus, midbrain, striatum, and thalamus) that represent the high, middle and low nicotinic receptor levels in the brain are presented in this study. Due to the small size of some brain regions (e.g., hippocampus), partial volume effects and activity spillover from neighboring regions may lead to some under or over estimation of radiotracer when using the software to extract the volumes-of-interest (VOI). The VOI of the whole brain was 473±3.39 mm^3^ (mean±SEM).

### Distribution volume ratio calculation

PMOD kinetic modeling software (Bruker, Zürich, Switzerland) was used to compute the distribution volume ratio (DVR) of the thalamus and midbrain of each mouse using Logan graphical analysis (Logan, Fowler et al. 1996) with the cerebellum used as the reference region. Using the concentration TACs generated from the VOIs from the VivoQuant brain atlas, a simplified reference tissue model (SRTM) was used to compute the *k*_2_’ value for each region. Then a Logan plot was created, and the slope used as the DVR.

### Statistical Analyses

Statistical analyses were performed using Prism software (GraphPad, La Jolla, CA, USA). All cellular data and all PET data were normally distributed. Continuous data were summarized as mean with standard errors with statistically significant differences assessed using t-test between two groups and one-way and two-way ANOVA among multiple groups. p<0.05 was considered statistically significant.

## Results

### Trapping of α4β2R PET ligands in α4β2R-containing intracellular acidic vesicles

Previously, we have found that the degree to which weak-base α4β2R ligands are trapped is dependent on both their pKa values and their affinity values for α4β2R binding sites (Govind, Vallejo et al. 2017) (see Table 1 for values). In order to estimate α4β2R affinity values for the PET ligands, Nifene and 2-FA (see structure in Fig 1 A), we performed competitive binding of the two ligands and nicotine using [^125^I]-Epb binding to α4β2Rs expressed in HEK cells. The Ki values from the fits to the data (Fig 1 B) were 7.1 x10^-11^ M for 2-FA, 3.1×10^-10^ for Nifene and 9.4 x10^-10^ for nicotine indicating that Nifene has a higher affinity for α4β2R binding sites than nicotine, and 2-FA a higher affinity than Nifene. These estimates (Table 1) of 2-FA and Nifene affinities for α4β2R binding sites together with their pKa values (Hansch, Corwin et al. 1995, Davila-Garcia, Musachio et al. 1997, Valette, Bottlaender et al. 1999, Brown, Chefer et al. 2004, Pichika, Easwaramoorthy et al. 2006, Lu, Chen et al. 2007, Gao, Kuwabara et al. 2008, Govind, Vallejo et al. 2017, Thompson, Metzger et al. 2017, Mukherjee, Lao et al. 2018) suggest that 2-FA and Nifene could become trapped in α4β2R-containing intracellular acidic vesicles similar to varenicline and epibatidine.

**Figure 1.**
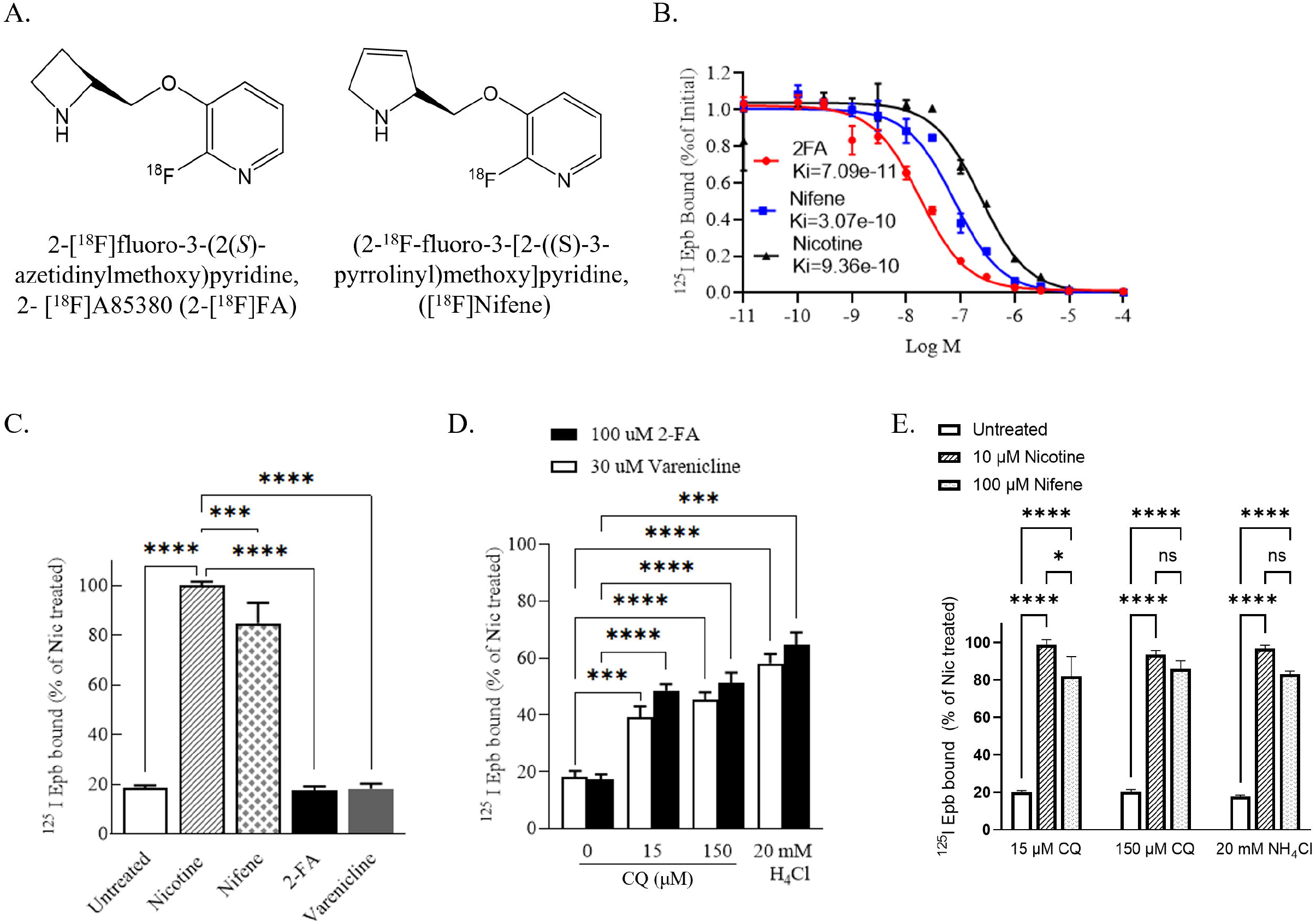
**A.** Chemical structures of 2-[^18^F]fluoro-3-(2(*S*)-azetidinylmethoxy)pyridine and (2-^18^F-fluoro-3-[2-((S)-3-pyrrolinyl)methoxy]pyridine. **B.** Competition of the ligands, nicotine (black triangle), Nifene (blue square) or 2-FA (red circle), for high affinity binding sites on α4β2Rs as assayed by ^125^I-epibatidine binding. Live α4β2R-expressing HEK cells were incubated with the indicated concentrations of nicotine, Nifene or 2-FA for 5 min prior to ^125^I-epibatidine binding. Data are plotted as the fraction of the initial ^125^I-epibatidine bound to intact cells in the absence any added ligands. Each point is the mean ± SEM. **C.** Trapping of 2-FA and not Nifene in intact α4β2R HEK cells. Cells were treated for 17 hours with Nicotine (10 μM), Nifene (100 μM), 2-FA (100 μM) or varenicline (30 μM) and ^125^I-epibatidine binding performed on live cells (n = 3). Specific epibatidine binding was represented as % of binding relative to nicotine treated cells. **D.** Loss of trapped 2-FA by disrupting intracellular pH gradient. α4β2R HEK cells were treated with 2-FA and varenicline for 17 hours. Cells were either washed with PBS or exposed to agents that raise pH in intact cells like chloroquine (15 μM or 150 μM) or NH4Cl (20 mM) for 10 minutes. ^125^I-epibatidine binding was performed and data plotted as in C. Each point is the mean ± SEM, n=3 for 2-FA and n=5 for varenicline. E. Disrupting intracellular pH gradient had no effects on untreated cultures or cultures treated with nicotine or Nifene. α4β2R HEK cells were untreated or treated with nicotine or Nifene for 17 hours and then treated with chloroquine (15 μM or 150 μM) or NH_4_Cl (20 mM) for 10 minutes as in C. In all the column graphs in (C, D and E): *p<0.05; **p<0.001; ***p<0.0001; ****p<0.00001 by one-way ANOVA with Bonferroni’s multiple comparison test.

To assay for the trapping of Nifene and 2-FA in α4β2R-containing intracellular acidic vesicles, we performed a different assay that also made use of [^125^I]-Epb. As described previously (Govind, Vallejo et al. 2017), the assay measures the degree to which a 17-hour exposure to the ligand increased the number of high-affinity α4β2R binding sites using [^125^I]-Epb binding, which is a measure of α4β2R upregulation by the ligand. If the assay is performed on live HEK cells expressing α4β2Rs, varenicline (Fig. 1C) and epibatidine (Govind, Vallejo et al. 2017) do not cause any significant upregulation compared to the 5-fold upregulation with nicotine exposure (Fig. 1C). However, if the assay is performed using membranes prepared from the cells in which intracellular acidic vesicles are not intact, α4β2R upregulation by varenicline and epibatidine is similar to that by nicotine (Govind, Vallejo et al. 2017). These data are consistent with varenicline and epibatidine being concentrated and trapped within the α4β2R-containing intracellular acidic vesicles, thereby preventing [^125^I]-Epb binding because varenicline and epibatidine are occupying the α4β2R binding sites in the vesicle lumen. Like varenicline, α4β2Rs were not upregulated by 2-FA for live HEK cells (Fig. 1C). In contrast, α4β2Rs were upregulated by Nifene almost to the same degree as upregulation by nicotine (Fig. 1C). These findings suggest that 2-FA is trapped within α4β2R-containing intracellular acidic vesicles similar to varenicline while Nifene is largely not trapped.

Upregulation by varenicline and epibatidine could be partially restored for live HEK cells by different treatments that dissipated the pH gradient across the vesicle membrane. These included the addition of a weak base that is not a α4β2R ligand, chloroquine (CQ) or ammonium chloride (NH_4_Cl). Consistent with 2-FA trapping within α4β2R-containing intracellular acidic vesicles, both chloroquine at 15 or 150 μM and ammonium chloride at 20 mM increased the effects of 2-FA and varenicline in terms of the upregulation of α4β2Rs in the HEK cells (Fig. 1D). In contrast, chloroquine and ammonium chloride, treatments that dissipated the pH gradient across the vesicle membrane, had no effect on upregulation by nicotine and Nifene consistent with a lack of trapping (Fig. 1E).

### Are α4β2R-containing intracellular acidic vesicles Golgi satellites?

We have recently characterized changes in the Golgi apparatus that occur with nicotine exposure and parallel upregulation of α4β2Rs (Govind, Jeyifous et al. 2021). With nicotine exposure, small organelles, Golgi satellites (GSats), form out of the endoplasmic reticulum (ER). GSat formation only occurs in response to nicotine when α4β2Rs are expressed in HEK cells or in neurons and the effects of nicotine are reversible. GSats had not been characterized previously because the usual Golgi markers used to image Golgi are Golgi structural proteins that are not found in GSats (Govind, Jeyifous et al. 2021). To examine whether the α4β2R-containing intracellular acidic vesicles are Golgi satellites we used the Golgi enzyme sialyltransferase Gal3 (St3) which label GSats in addition to the Golgi apparatus (GA).

In Fig. 2, we imaged GSats by expressing a GFP-tagged version of St3 in cultured rat cortical neurons together with α4 and HA-tagged β2 subunits. St3-GFP labels the GA found in the soma of untreated neurons (Fig. 2, -Nicotine panels) or neurons that have been exposed to nicotine for 17 hours (Fig. 2, +Nicotine panels), which increases the numbers of GSats (Govind, Jeyifous et al. 2021). St3-GFP also labels smaller GSat structures that extend out into the neuronal processes. Two different examples of neurons are displayed in Fig. 2A and 2B both expressing St3-GFP (green), but with the α4β2Rs labeled using different probes. In Fig. 2A, the α4β2Rs were labeled with fluorescent anti-β2 antibody that specifically labels all β2 subunits assembled and unassembled (Walsh, Roh et al. 2018), which includes transfected and endogenous subunits. The labeled β2 subunits are found largely in the ER where β2 subunits are synthesized and assemble with α4 into α4β2Rs (Sallette, Pons et al. 2005). The probe also stains the GA but appears to have little overlap with the smaller St3 puncta, which are the GSats. This apparent lack of overlap results from the very bright staining of the β2 subunits in the ER relative to β2 subunit staining of the Golgi satellites. The presence of β2 subunits in GSats is observed when we increase the resolution of the images in the soma (see insets in Fig. 2A) especially in the +Nicotine image. The β2 subunit staining of the GSats in even better observed in dendrites (Fig. 2C(i)) where the levels of ER are much smaller, and consequently, staining of β2 subunits in ER is much weaker.

**Figure 2.**
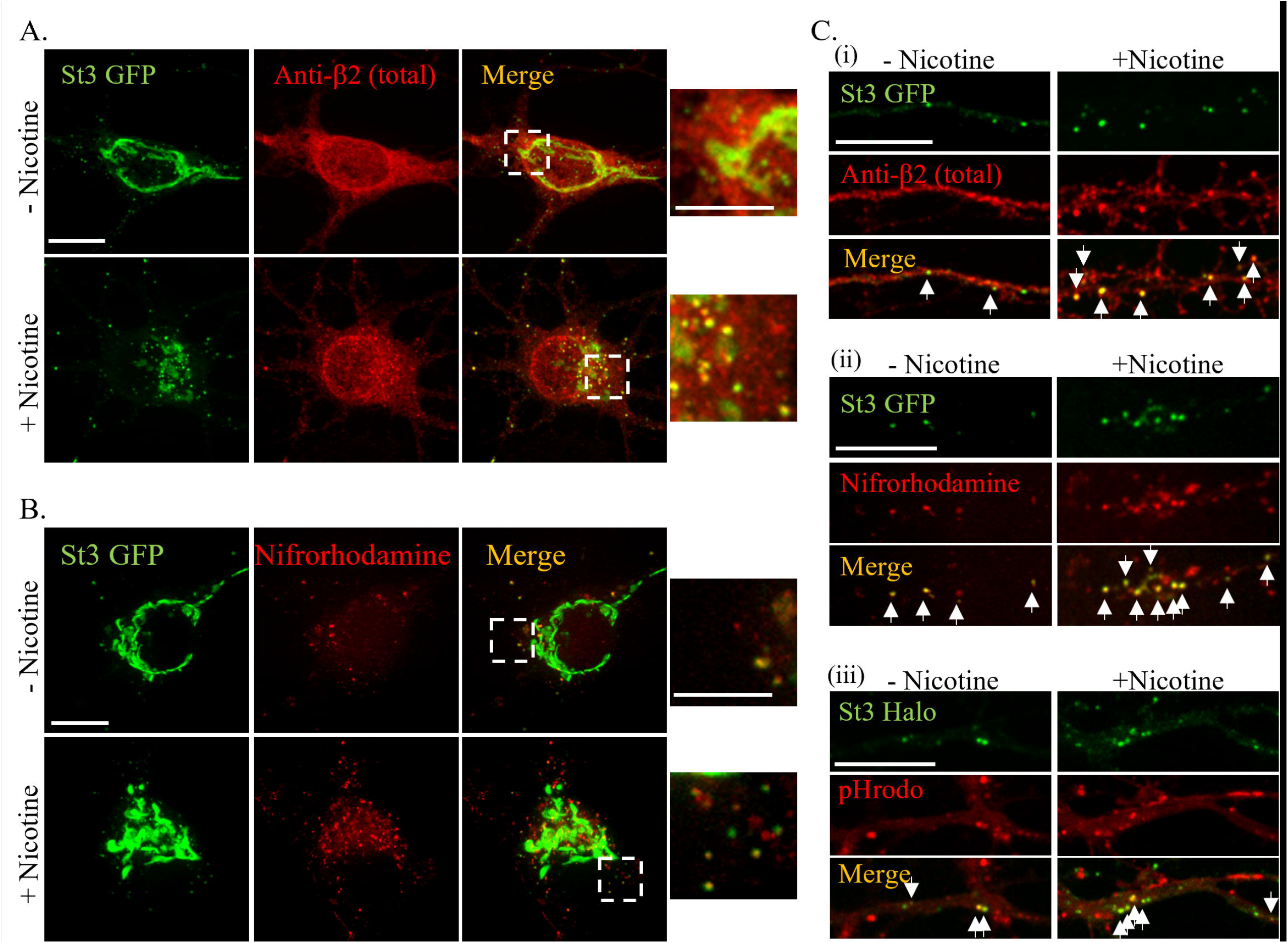
Nifrorhodamine labeling of GSats in primary cortical cultures expressing α4β2Rs. **A.** Cultured neurons (DIV 10) were expressing the GSat marker St3-GFP (green) and α4 and β2HA subunits. After 24 hours, cultures were treated with 1 μM nicotine for 17 hours (+Nicotine) or untreated (-Nicotine). Neurons were fixed, permeabilized and stained for all β2 subunits with monoclonal anti-β2 antibody, mAb12H (red; secondary antibody anti-mouse Alexa 568). Scale bar, 10 μm. Inset scale bar, 5 μm. **B.** High-affinity α4β2R binding sites were labeled with Nifrorhodamine in primary cortical cultures expressing α4β2Rs. Cultured neurons were transfected and treated with nicotine as in A. Cultures were labeled with 25 μM Nifrorhodamine (red) for 30 minutes and imaged live. Scale bar, 10 μm. Inset scale bar, 5 μm. Inset scale bar. **C.** Dendritic GSats contain high-affinity α4β2R binding sites and are acidic. (i). β2 subunit labeling of dendrites of neurons expressing St3-GFP (green) and α4 and β2-HA subunits (anti-β2-mAb12H, red) after permeabilization and fixation as in A. (ii). Live labeling of Nifrorhodamine (red) with St3-GFP (green) as in B. (iii). Live labeling with pH-sensitive dye, pHrodo (red), with St3-Halo expression (green). Arrows mark Golgi satellites with co-labeling for β2-HA subunits, Nifrorhodamine or pHrodo. Scale bar, 10 μm.

In Fig. 2B, α4β2Rs were labeled using a membrane-permeable, fluorescently-labelled analogue of the 2-FA ligand, Nifrorhodamine, that binds to α4β2R high-affinity binding sites. In order to obtain a fluorescent probe for α4β2Rs, significant modification in the structure was necessary due to the lack of functionality of 2-FA and Nifene for fluorophore labeling (see (Samra, Intskirveli et al. 2018) for details). The modifications necessary to add rhodamine to 2-FA resulted in an analogue closer to Nifrolidine, which is a similar α4β2R ligand used for PET imaging (Chattopadhyay, Xue et al. 2005). Surprisingly, Nifrorhodamine labeling does not colocalize with β2 subunit staining in the ER or GA in the soma. Instead, Nifrorhodamine labeling co-localizes with the GSats labeled with St3-GFP in the soma (Fig. 2B) and dendrites (Fig. 2C(ii)). These findings suggest that α4β2R high-affinity binding sites form in GSats and not during assembly in the ER. Shown in the right panels of Fig. 2C (iii) are dendrites expressing St3-GFP and labeled with pHrodo, a fluorescent probe that fluoresces at pH values found in acidic vesicles. pHrodo fluorescence overlays with a significant number of the St3-positive puncta as well as other acidic organelles consistent with mature Golgi satellites being acidic vesicles.

### PET studies with 2-[^18^F]FA and [^18^F]Nifene

We measured the binding kinetics of α4β2R ligands *in vivo*, using 2-[^18^F]FA and [^18^F]Nifene for brain PET imaging in mice. Figure 3A displays typical images of 2-[^18^F]FA and [^18^F]Nifene binding at coronal, sagittal, and transverse sections of the brain in WT mice. Whole brain uptake of [^18^F]Nifene was significantly greater than 2-[^18^F]FA as seen in Fig-3A. The TACs of the whole brain from the WT mice for 2-[^18^F]FA and [^18^F]Nifene are shown in Figure 3B. [^18^F]Nifene demonstrated fast kinetics peaking at ~10 minutes with a SUV of ~1.6 after tracer injection. In contrast the binding of 2-[^18^F]FA was slower peaking at ~50 minutes with a SUV of ~0.6. The significant difference in brain kinetics between the two radiotracers is seen in the inset of Fig-3B where whole brain [^18^F]Nifene reduces to below 50% levels within 40 minutes whereas 2-[^18^F]FA remained at ~70% of the peak at the end of 3 hour imaging time.

**Figure 3.**
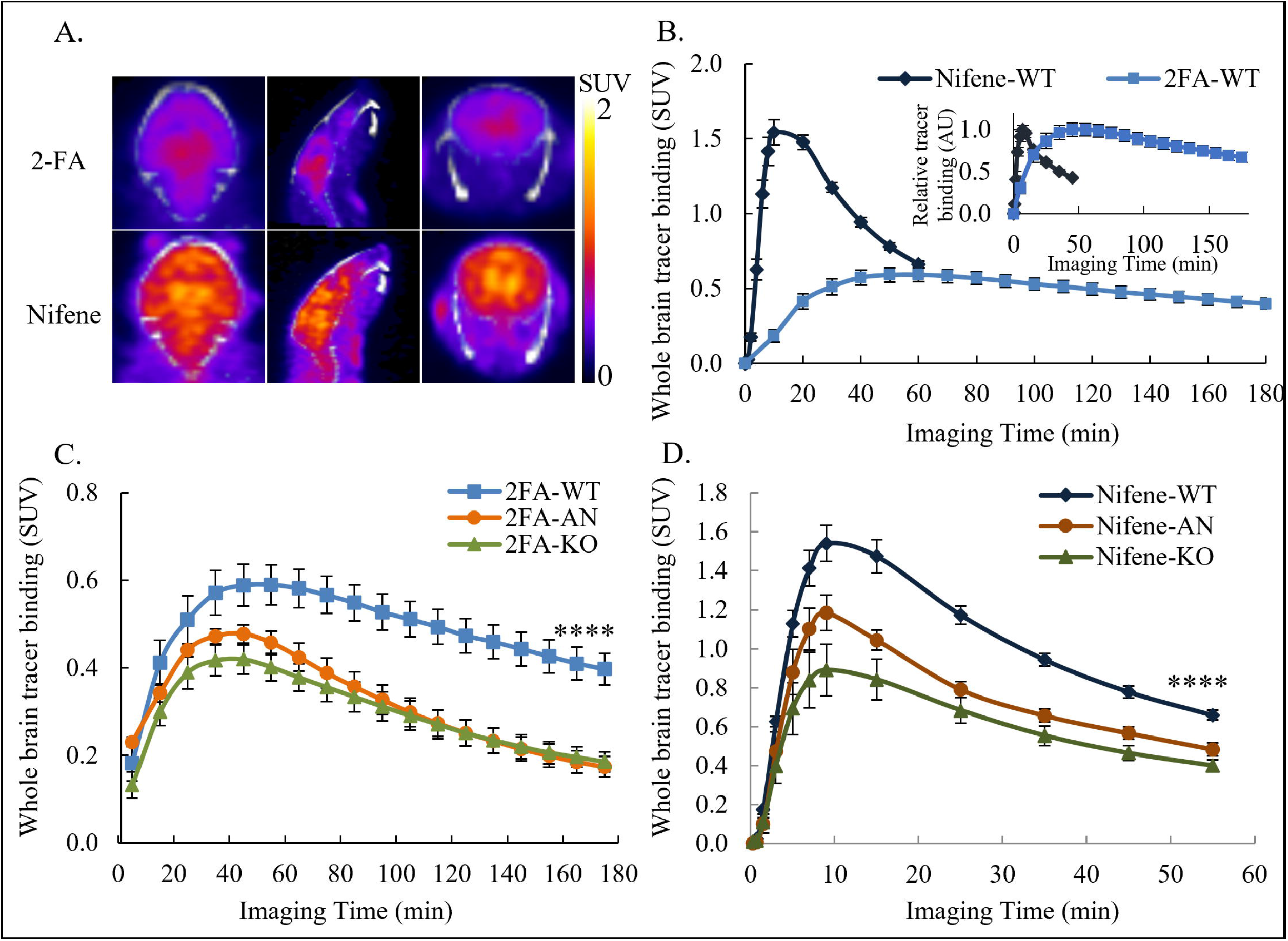
Binding and disassociation of 2-[^18^F]FA to α4β2Rs is slower than [^18^F]Nifene. **A.** Brain images of 2-[^18^F]FA and [^18^F]Nifene binding at coronal, sagittal, and transverse sections in WT mice. **B.** Whole brain time active curves show [^18^F]Nifene has faster binding kinetics compared to 2-[^18^F]FA. The insert compares both tracers after signal normalization to their highest binding respectively. **C & D.** Acute nicotine administration and β2-subunit knockout diminish the binding of 2-[^18^F]FA & [^18^F]Nifene.

The whole brain retention for both 2-[^18^F]FA and [^18^F]Nifene was significantly decreased in the β2 KO mice. This is similar to our previous observations of a lack of [^18^F]Nifene binding in β2 KO mice (Bieszczad, Kant et al. 2012). Similarly, 2-[^18^F]FA cleared from the β2 KO mice brain more rapidly, with SUV values reduced by approximately 50% of WT animals at 3 hrs post-injection (Fig-3C). Reductions in whole brain SUVs of [^18^F]Nifene were also approximately 50% in the β2 KO mice compared to WT animals (Fig-3D).

Acute nicotine challenge in WT animals affected the whole brain uptake and retention of both 2-[^18^F]FA and [^18^F]Nifene. The initial higher uptake of both 2-[^18^F]FA and [^18^F]Nifene compared to the KO mice may be due to the higher free fraction caused by the nicotine challenge. In the case of 2-[^18^F]FA, acute nicotine brought the SUV levels at the end of the 3 hr scan similar to the β2 KO mice brain (Fig-3C). Retention of [^18^F]Nifene after acute nicotine was similarly reduced with clearance profile similar to that found in the β2 KO mice brain (Fig-3D).

### [^18^F]Nifene Binding in Brain Regions

Regional brain distribution of [^18^F]Nifene binding was analyzed in the thalamus, midbrain, hippocampus, striatum, cortex and cerebellum. In wild type mice, [^18^F]Nifene showed the highest SUV in the thalamus and midbrain and the lowest SUV in the cerebellum throughout the imaging duration (Fig 4A). Time activity curves in the various regions followed a similar profile in all brain regions. Levels of [^18^F]Nifene binding in the β2 KO mice brain were reduced to the level of cerebellum in all regions (Fig 4B). In the case of acute nicotine administration initial levels of uptake were higher than β2 KO mice (Fig 4C), which may be due to higher free fraction of [^18^F]Nifene, but subsequently retention of [^18^F]Nifene was not observed in receptor rich regions, unlike the WT without nicotine (Fig 4A).

**Figure 4.**
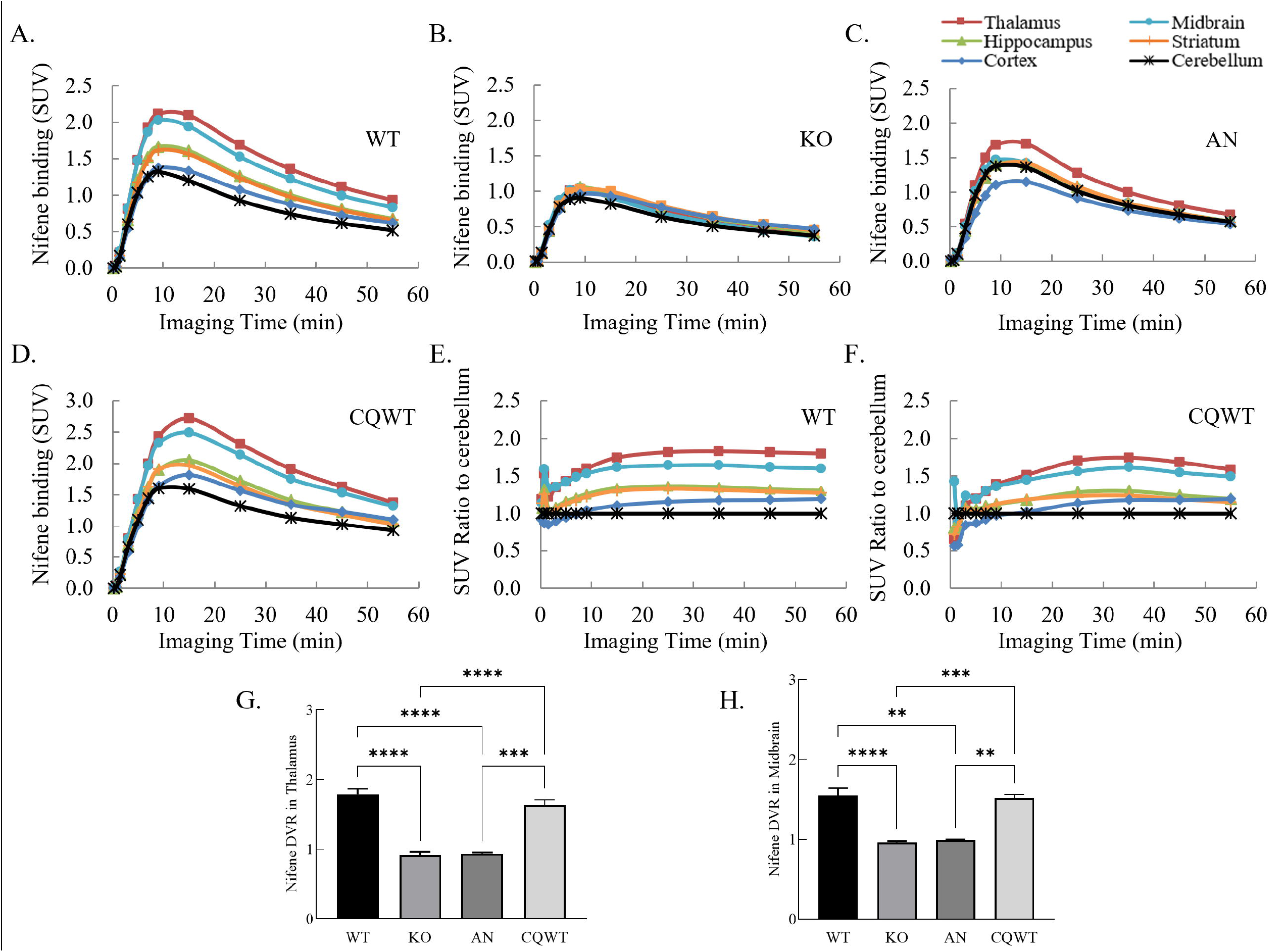
TACs of 6 brain regions shows that the thalamus and midbrain have the highest α4β2R ligand binding while the cerebellum represents non-specific binding. **A.** TACs of [^18^F]Nifene in 6 selected brain regions of WT mice. **B.** TACs of [^18^F]Nifene in KO mice. **C.** TACs of [^18^F]Nifene in AN mice. **D.** TACs of [^18^F]Nifene in chloroquine (CQ) treated WT mice. **E & F.** Binding ratios of each brain region to the cerebellum from WT mice and mice treated with CQ. **G & H.** Nifene DVR in the thalamus and midbrain of WT, KO, AN and CQWT mice. KO and AN mice show reduced specific binding compared to WT mice. CQWT mice show no difference in DVR to WT mice.

Fig 4D displays the TACs for CQWT mice, and compared to the WT mice (Fig 4A), no difference in [^18^F]Nifene uptake was noticed following CQ treatment. This is further illustrated by the SUV ratio to cerebellum curves for WT mice (Fig 4E) and CQWT mice (Fig 4F) showing similar levels of [^18^F]Nifene specific binding between groups.

Using the cerebellum as the reference region, DVR of [^18^F]Nifene across all groups of mice are shown for the thalamus (Fig 4G) and midbrain (Fig 4H). The DVR values for β2 KO mice and acute nicotine treated mice were <1. The wild type mice had DVRs approaching 2 in the thalamus and over 1.6 in the midbrain. Differences between the WT and β2 KO mice were highly significant, as well as the difference between WT and acute nicotine-treated animals. The similar DVR levels of β2 KO mice and acute nicotine-treated animals confirms the apparent lack of receptor-mediated specific binding. No significant difference was observed between [^18^F]Nifene DVR in the WT and CQWT mice, indicating that CQ treatment does not affect the binding of [^18^F]Nifene *in vivo*, consistent with our observations *in vitro* (Fig. 1).

### 2-[^18^F]FA Binding in Brain Regions

In WT mice, 2-[^18^F]FA showed the highest SUV (~0.7) in the thalamus and midbrain and the lowest SUV in the cerebellum (~0.2-0.3) at the end of the imaging session (Fig 5A). Other regions such as the striatum and cortex exhibited moderate levels of 2-[^18^F]FA binding. Compared to [^18^F]Nifene (Fig. 4A), 2-[^18^F]FA SUVs did not reach a plateau within the three-hour scanning period. However, given that the residence time of varenicline in the brain is 4-5 days, the kinetics of 2-[^18^F]FA appear consistent to the slow kinetics of varenicline. All brain regions in the β2 KO mice exhibited a similar uptake and clearance profile with SUVs similar to cerebellum of ~0.2-0.3 (Fig 5B). Similarly, acute nicotine was also able to block retention of 2-[^18^F]FA binding in various brain regions with SUVs similar to cerebellum of ~0.3 (Fig 5C).

**Figure 5.**
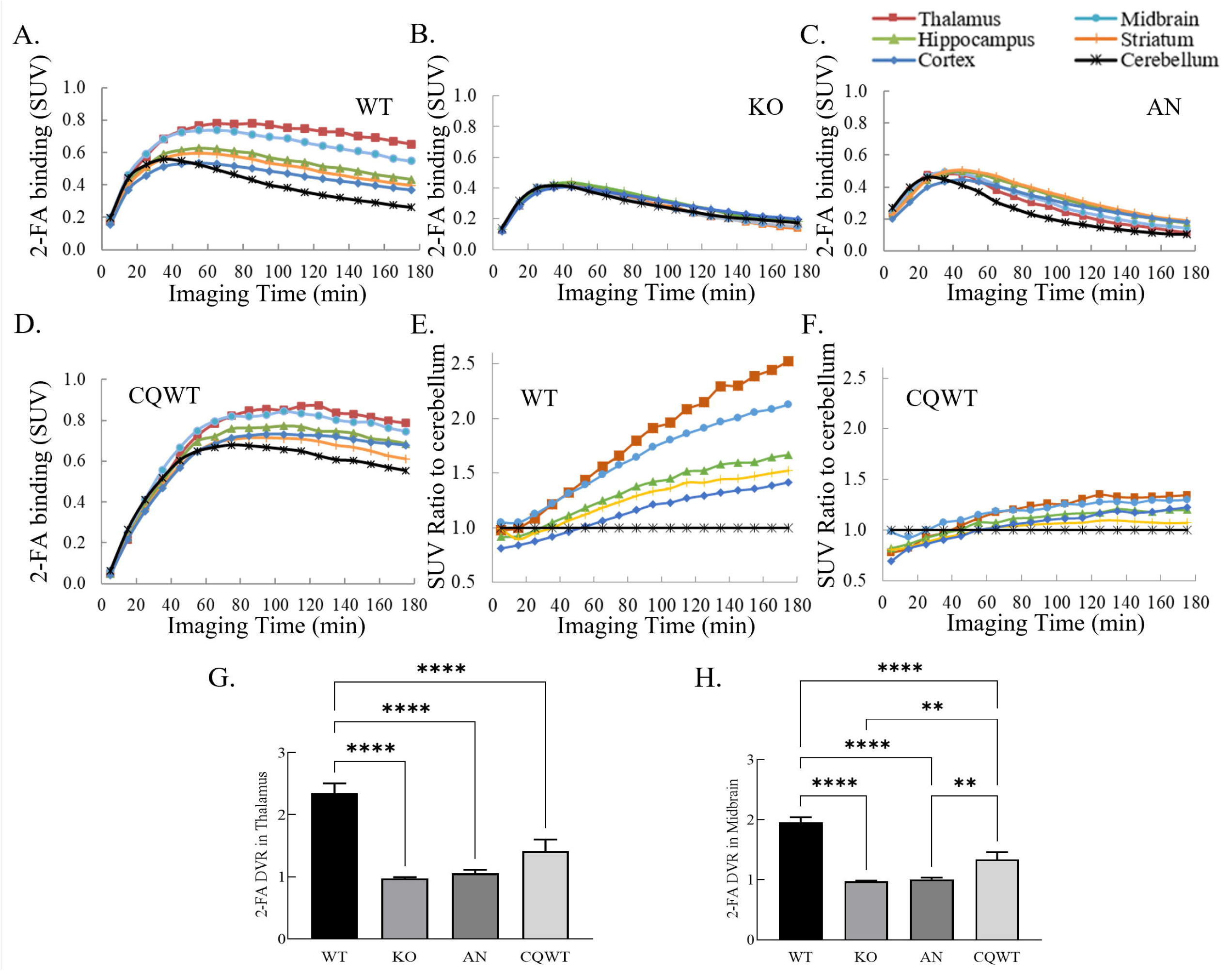
*In vivo* CQ treatment decreases the binding of 2-[^18^F]FA and slows its disassociation in 6 selected brain regions. **A.** TACs of 2-[^18^F]FA in 6 selected brain regions from WT mice. **B.** TACs of 2-[^18^F]FA in KO mice. **C.** TACs of 2-[^18^F]FA in AN mice. **D.** TACs of 2-[^18^F]FA in chloroquine (CQ) treated WT mice. **E & F.** Binding ratios of each brain region to the cerebellum from WT mice and mice treated with CQ. **G & H.** 2-[^18^F]FA DVR in the thalamus and midbrain of WT, KO, AN and CQWT mice. KO, AN and CQWT mice have reduced specific binding compared to WT mice.

Because the cerebellum had the lowest levels of 2-[^18^F]FA, DVR in other regions used the cerebellum as the reference region. The thalamus had the highest DVR (>2) in WT mice followed by the midbrain DVR (~2) (Fig 5G, H). The DVR values for β2 KO mice and acute nicotine-treated mice were <1. Differences between the WT and β2 KO mice were highly significant, as well as the difference between WT and acute nicotine-treated animals. The similar DVR levels of β2 KO mice and acute nicotine-treated animals confirms the apparent lack of receptor-mediated specific binding.

To further test whether the mechanism causing 2-[^18^F]FA slow kinetics, mice were treated with CQ, which as discussed previously, dissipates the pH gradient across acidic organelles and reduced trapping *in vitro* (Fig. 1D; (Govind, Vallejo et al. 2017)). Previous studies had established that the maximum amount of CQ that can be injected into mice resulted in blood level of ~15 μM CQ for 3 days before PET imaging with 2-[^18^F]FA. Increased uptake of 2-[^18^F]FA was observed in several brain regions post CQ treatment. This increase in 2-[^18^F]FA uptake may reflect an effect on the ability of 2-[^18^F]FA to sequester in intracellular acidic vesicles throughout the brain, which would increase levels of free 2-[^18^F]FA. Compared to the regional binding difference in WT mice (Fig 5A), CQ reduced the regional differences of 2-[^18^F]FA binding although the nonspecific 2-[^18^F]FA binding increased in CQWT mice (Fig 5D). This reduction in specific binding to α4β2Rs is more evident in the binding ratios of different regions to cerebellum (Fig 5E and 5F). Total activity uptake of 2-[^18^F]FA in thalamus of the WT animals was approximately 2.5 fold over that in cerebellum at the end of 3 hour imaging, while it was less than 1.5 fold in CQ treated animals. Similarly, uptake of 2-[^18^F]FA in other brain regions were also reduced. The decrease in thalamic and midbrain DVR of CQ treated mice were statistically significant (Fig 5G and 5H). Thus, the results demonstrate that CQ treatment of the mice significantly reduced the specific 2-[^18^F]FA signal in the thalamus and midbrain consistent with it reducing 2-FA trapping as observed *in vitro* (Fig. 1D).

## Discussion

Despite the known adverse consequences associated with smoking, smoking cessation is difficult. Even with the most effective smoking cessation reagent, varenicline, quitting rates reach only ~50% (Agboola, Coleman et al. 2015). To rationally design better smoking cessation drugs, it is critical to understand the molecular and cellular mechanisms governing how drugs dampen the effects of nicotine on the brain’s reward pathway. Varenicline was designed as a high-affinity, partial agonist that mediates anti-smoking effects through acute, rapid interactions with α4β2Rs that reduce the full-agonist effects of nicotine. Our lab has discovered additional long-term effects of varenicline in which varenicline is trapped in intracellular acidic vesicles and maintain brain levels throughout the day, importantly, after nicotine levels decline (Govind, Vallejo et al. 2017). We demonstrated that the slow release of trapped varenicline reduced the functionally upregulated α4β2R currents, which should contribute to varenicline effects on smokers (Govind, Vallejo et al. 2017).

The ‘trapping hypothesis’ has only been tested using *in vitro* systems. However, the long-term trapping of varenicline in GSats in the human brain explains why the rate of varenicline disposition is much slower (4-5 days) compared to that of untrapped and rapidly metabolized nicotine (1-2 hours; (Govind, Vallejo et al. 2017). Here, we extended our analysis *in vivo* using PET imaging in mice. Based on the pKa and α4β2R affinity values (Table 1), we predicted that the α4β2R PET ligand, 2-FA, is trapped in acidic vesicles similar to varenicline, and Nifene would have significantly less trapping. 125I-epibatidine binding *in vitro* to measure for reduction in α4β2R upregulation as a measure of trapping confirmed that 2-FA is trapped in acidic vesicles similar to varenicline and Nifene displayed little trapping similar to nicotine (Fig. 1C).

Radioligands 2-[^18^F]FA and [^18^F]Nifene were chosen based on their pKa and α4β2R binding affinity values with those of 2-FA similar to varenicline and Nifene similar to nicotine (see Table 1). Additionally, the previously unexplained slow (2-FA) and fast (Nifene) PET kinetics of nicotinic receptor binding *in vivo* are consistent with our trapping hypothesis. We successfully employed both 2-[^18^F]FA and [^18^F]Nifene in PET/CT mouse brain imaging of α4β2Rs, making this study the first to use 2-[^18^F]FA in mouse models. As observed for studies in human and other animal models, the 2-[^18^F]FA uptake in the whole brain is much slower than that of [^18^F]Nifene, peaking at ~50 min vs ~10 min after the intraperitoneal tracer administration (Fig. 3 B-D). This difference also holds true for the clearance of the two ligands from the brain (Fig. 3B). We observed statistically significant reduction of ligand binding in the whole brains of mice after acute nicotine injections and in the β2-subunit knockout mice, demonstrating that most specific binding sites are found on β2-containing receptors.

Structural differences between 2-FA and Nifene can explain differences in the rates of entry into the brain and initial differences in binding levels (Fig. 3B-D). Both ligands have similar features: a 2-fluoropyridine group, a methoxyether linkage at the 3-position of the pyridine ring, and a secondary nitrogen in the 4-membered azetidine ring in 2-FA and 5-membered 3,4-dehydropyrrolidine ring in Nifene (Fig. 1A). However, the saturated 4-membered ring in 2-FA versus an unsaturated 5-membered ring in Nifene makes 2-FA more basic (pKa >10), while Nifene’s unsaturated ring, which can delocalize electron density away from the nitrogen, is less basic (pKa ~9). This difference in pKa renders 2-FA more hydrophilic relative to Nifene. The lower lipophilicity and higher pKa of 2-FA make it less permeable into the brain, due to a lower proportion of available uncharged species to cross the blood brain barrier (BBB), whereas Nifene has a higher proportion of uncharged species which freely crosses into the brain. This difference in permeability is reflected in Fig 3B-D with a high initial uptake of Nifene compared to 2-FA. The higher affinity of 2-FA and a high nonspecific binding component due to the higher pKa also favors binding-rebinding of 2-FA at the receptor sites, further slowing down the dissociation kinetics. Nifene, with lower affinity and lower pKa can more rapidly equilibrate and dissociate from the receptor sites, thus having a faster clearance rate.

Further analyses showed that two ligands displayed identical regional uptake with the highest uptake within the thalamus and midbrain (Fig. 3A), consistent with PET imaging studies in humans and other animal models (Hillmer, Wooten et al. 2011, Hillmer, Wooten et al. 2012, Hillmer, Wooten et al. 2013, Mukherjee, Lao et al. 2018). Regional binding differences were diminished in AN mice and completely abolished in KO mice (Fig. 3C-D). These findings demonstrate that the ligand uptake in the cerebellum represents mostly non-displaceable ligand signal. However, the WT mice displayed higher SUVs in the cerebellum compared to AN and KO mice for both 2-[^18^F]FA and [^18^F]Nifene, indicating the presence of some ligand specific binding. This cerebellum specific binding signal is not unique to mice, as studies in rats using 2-[^18^F]FA PET found a nicotine displaceable signal component in this region (Vaupel, Stein et al. 2007). A study using [^18^F]Nifene in rats also noticed significant ligand displacement in the cerebellum following administration of both lobeline and (-)nicotine, indicating the presence of specific binding (Hillmer, Wooten et al. 2013). Due to the presence of this specific binding signal, DVR calculations using a cerebellum reference region will underestimate the binding potential. Because our study design takes advantage of AN and KO mice whose cerebellar specific binding was completely abolished, only DVR values of the WT mice underestimate the true binding potential. Despite the underestimation, the WT mice displayed significantly higher DVR compared to all other groups, indicating that our choice of reference region was suitable for this application. The above-18 mentioned studies suggest the corpus collosum may be a suitable reference tissue for 2-[^18^F]FA and [^18^F]Nifene PET studies. The corpus collosum was explored as a reference tissue for the current study, but displayed higher SUVs compared to the cerebellum in WT mice and displayed higher SUVs in WT mice compared to KO and AN mice. In addition, due to the small volume of the corpus collosum, the signal from this region was likely confounded by partial volume effects. Future studies should consider alternative reference regions or measurement of arterial blood for improved PET quantification.

Chloroquine, which is a BBB permeable weak base and not a nicotinic ligand, has been used widely to reduce the pH gradient across intracellular acidic compartments. The decrease of the pH gradient across intracellular acidic vesicles has been directly measured in cultured cells (Dong, Song et al. 2017). We found that inclusion of chloroquine in the medium at concentrations of 15-150 μM or NH_4_Cl at 20 mM significantly reduced trapping of varenicline and 2-FA in cultured cells (Fig. 1D). We assume the reduction in trapping is caused by chloroquine entry in the α4β2R-containing Golgi satellite vesicles reducing the pH gradient through their protonation without interfering with nicotinic ligand binding to α4β2Rs so that weak base nicotinic ligand is less trapped. Comparing the 2-[^18^F]FA SUV TACs between the CQ mice and the untreated WT mice revealed that chloroquine resulted in uptake of the ligand in the target regions to more closely resemble the cerebellum (evident in Fig. 5 E-F), indicating reduced specific binding in these regions and significantly faster 2-[^18^F]FA kinetics. Under the presence of chloroquine, regional brain uptake was higher than in the cerebellum, indicative of the presence of specific ligand binding. These results are consistent with 2-[^18^F]FA being trapped within intracellular acidic vesicles, the Golgi satellites, *in vivo* in the mouse brains. In contrast, CQ treatment prior to Nifene imaging did not result in changes of Nifene binding (Fig. 4 D-H), suggesting that Nifene is not trapped in Golgi satellites *in vivo*.

Of interest is the use of dynamic PET data to measure the efflux rate of 2-FA from the acidic vesicles to compare with the *in vitro* findings. When generating the Logan plots, the tissue-to-plasma efflux rate (*k*_2_ ’) was determined using SRTM with a cerebellum reference tissue. For 2-FA, the efflux rate was determined as 0.033±0.0030 min^-1^ (mean±SEM). Compared to the efflux rate determined for Nifene (0.45±0.084 min^-1^), 2-FA reveals much slower washout from the brain. Future studies will explore measurement of the arterial blood curve and use of an image derived input function in different compartment models to estimate these rate constants, where the value of *k_4_* is likely to be a more accurate measure of the efflux of ligand from the intracellular acidic vesicles.

Taken together, our results are consistent with 2-FA and Nifene interacting with α4β2Rs by different mechanisms. Because Nifene does not get trapped significantly in the α4β2R-containing GSats, its exit from neurons expressing α4β2Rs is only delayed by the binding and unbinding to the receptors and is a measure of the total α4β2R population, both cell-surface and in intracellular receptors. 2-FA binds to α4β2Rs in the Golgi satellites and becomes trapped for many hours. Because its binding and unbinding to receptors outside the Golgi satellites is so much faster than its exit rate from the Golgi satellites, its PET signal in the brain is a measure of only Golgi satellite pools of receptors. In short, the two ligands measure separate α4β2R features. In this study, we have combined *in vitro* cellular assays with *in vivo* PET and new insights into the changes that occur during nicotine addiction have emerged that could help design more effective anti-smoking drugs. Specifically, our findings suggest that design elements, such as pKa and receptor affinity that regulate trapping in the Golgi satellites, need to be explored in terms of drug design. The same drug design strategy can be applied to other neurological conditions in which nicotinic receptors are implicated such as dementia (Picciotto, Brunzell et al. 2002), depression (Picciotto, Brunzell et al. 2002) and attention deficits (Faraone and Biederman 1998).

## Acknowledgements

This study was supported in part by the NIH grant R01 DA044760-01 to J.M., C.C. and W.N.G., RF1 AG029479 to J.M., and T32 DA043469 to M.Z. The authors acknowledge the assistance from the Integrative Small Animal Imaging Research Resources (iSAIRR) supported in part by the NIH grant P30 CA14500 and S10 OD025265, and from the Cyclotron Facility of the University of Chicago.

